# Tracking of *Anopheles stephensi* in Ethiopia using mitochondrial DNA reveals pattern of spread

**DOI:** 10.1101/2021.04.07.437873

**Authors:** Tamar E. Carter, Solomon Yared, Dejene Getachew, Joseph Spear, Sae Hee Choi, Jeanne N. Samake, Peter Mumba, Dereje Dengela, Gedeon Yohannes, Sheleme Chibsa, Matthew Murphy, Gunawardena Dissanayake, Cecilia Flately, Karen Lopez, Daniel Janies, Sarah Zohdy, Seth R. Irish, Meshesha Balkew

## Abstract

The recent detection of the South Asian malaria vector *Anopheles stephensi* in the Horn of Africa (HOA) raises concerns about the impact of this mosquito on malaria transmission in the region. The mode and history of introduction is important for predicting the likelihood of continued introduction and future spread. Analysis of *An. stephensi* genetic diversity and population structure can provide insight into the history of the mosquito in the HOA. We investigated genetic diversity of *An. stephensi* in eastern Ethiopia where detection suggests a range expansion to this region to understand the history of this invasive population. We sequenced the cytochrome oxidase subunit I (*COI*) and cytochrome B gene (*CytB*) in 187 *An. stephensi* collected from 10 sites in Ethiopia in 2018. Phylogenetic analyses using a maximum-likelihood approach and minimum spanning network were conducted for Ethiopian sequences. Molecular identification of bloodmeal sources was also performed using universal vertebrate *CytB* sequencing. Six *COI-CytB* haplotypes were observed based on five segregating sites, with the highest number of haplotypes in the northeastern sites (Semera, Bati, and Gewana towns) relative to the southeastern sites (Kebridehar, Godey, and Degehabur) in eastern Ethiopia. In the phylogenetic and network analysis, we observed population differentiation based on the distribution of the haplotypes across the northeastern and central sites (Erer Gota, Dire Dawa, and Awash Sebat Kilo) compared to the southeastern sites and evidence of a South Asian origin of the HOA *An. stephensi* lineages. The presence of the putative South Asian haplotype of origin at sites closest to Ethiopia’s northeastern borders support route of introductions into Ethiopia from the northeast. Finally, molecular bloodmeal analysis revealed evidence of feeding on bovines, goats, dogs, and humans, as well as evidence of multiple (mixed) blood meals. In conclusion, we find support for the hypothesis for the recent expansion of *An. stephensi* into southeastern Ethiopia with multiple introductions. We also find evidence that supports the hypothesis that HOA *An. stephensi* populations originate from South Asia rather than the Arabian Peninsula. The evidence of both zoophagic and anthropophagic feeding support the potential for livestock movement to play a role in vector spread in this region.

## BACKGROUND

Malaria remains one of the leading global health concerns with over 229 million cases reported yearly ^1^. Efforts to prevent the transmission of malaria often involve controlling the mosquito vectors (*Anopheles*) that transmit the malaria parasite, *Plasmodium* spp. Previous studies have shown that the movement of *Plasmodium* strains from one region to another can have serious population health consequences such as the emergence of antimalarial resistance 2 or malaria epidemics in naïve or low transmission areas ^3^. Investigations into the long-distance movement of these new *Plasmodium* strains has typically centered on the movement of their human hosts as carriers for these strains, especially on asymptomatic individuals who act as reservoirs for *Plasmodium* transmission (for review, see Bousema et al. ^4^). With the growing evidence that vectors can move long distances ^5^ and the correlation of new vector introductions with increased disease burden in some instances ^6–8^, it is critical to investigate the movement of vector populations in malaria endemic regions in order to understand the introduction of new parasite strains and evaluate the malaria risk of a particular region.

The Horn of Africa (HOA), comprising Djibouti, Eritrea, Ethiopia, and Somalia, is classified as a malaria endemic area, with the distinction compared to most other areas in Africa of having substantial transmission of both *P. falciparum* and *P. vivax* ^1^. In 2012, an urban malaria vector mosquito, *Anopheles stephensi,* which is common in South Asia, was detected in the HOA for the first time in Djibouti in 2012 ^9,10^ and then in 2016 in east Ethiopia ^11^. Previously, this species range was believed to be limited to South Asia and the Middle East, including the eastern Arabian Peninsula ^12^. Like *An. arabiensis*, the major malaria vector in the HOA, *An. stephensi* is known to transmit both major malaria parasite species, *P. falciparum* and *P. vivax,* and could threaten recent progress in reducing the prevalence of malaria in that area. The first detection of *An. stephensi* in Africa was in Djibouti in 2012 ^9,10^ and then in 2016 in east Ethiopia ^11^. After additional surveillance, *An. stephensi* was also detected in the Republic of Sudan and Somalia (Vector Threat Map, WHO, 2019) and revealed to have a broader distribution in Ethiopia ^13^ and Djibouti 10 suggesting that *An. stephensi* may spread to other parts of Africa. With molecular evidence that various *Plasmodium* strains evolve to be compatible with specific *Anopheles* species, it was unclear whether *An. stephensi* could transmit local HOA *Plasmodium* strains. There is now evidence that the HOA *An. stephensi* can transmit the local strains of *P. falciparum* and *P. vivax* in Djibouti ^10^ and Ethiopia ^15^, generating concern for the role *An. stephensi* may play in increasing malaria transmission.

The expansion of *An. stephensi* in the HOA complicates malaria control in the HOA in several ways. Until now, vector control strategies have centered on the detection and elimination of known major vectors like *An. arabiensis* which exhibit different breeding, feeding, and resting behaviors when compared to *An. stephensi*. Also, there is evidence of diverse insecticide resistance mechanisms among species of *Anopheles* ^16–18^. Given these differences, vector control approaches are in the process of being modified to account for the presence of *An. stephensi* in the HOA. The potential threat to progress in controlling malaria led to the World Health Organization (WHO) posting a “vector alert” in 2019, characterizing *An. stephensi* a “major potential threat”, and calling for enhanced surveillance in Africa ^19^.

In order to better assess the potential impact of *An. stephensi* on malaria in the HOA and the rest of the continent, more information is needed on the pattern of spread of *An. stephensi*. Analysis of the genetic variation of *An. stephensi* populations can provide crucial information about the geographic extent of the species, and the frequency of introductions. Phylogeographic and population genetic analyses of the structure of variation and relationships between newly detected *An. stephensi* and the *An. stephensi* from established populations outside of Africa can provide insight into the history and evolution of this population. Initial analysis of the mitochondrial *COI* (cytochrome c oxidase subunit I) generated from *An. stephensi* in Kebridehar in east Ethiopia and *An. stephensi* sequence data from collections in South Asia, the Middle East, and the Arabian Peninsula taken from GenBank ^11^ identified a single haplotype, that was different from Djibouti sequences but identical to one *An. stephensi* sequence from Pakistan. Since the identification of *An. stephensi* in Ethiopia, this vector has been detected in ten additional sites in east Ethiopia as far north as Semera and as far south as Godey ^13^. Additional data from these sites are needed to provide a more complete picture of the history of *An. stephensi* in Ethiopia. In addition to phylogeographic history, more information on factors that facilitate *An. stephensi* spread is needed. One potential factor of interest is the frequent movement of livestock in the regions where *An. stephensi* has been detected. Pastoralism is a common practice in Ethiopia and involves moving cattle, goats, sheep, and camels hundreds of kilometers every season ^20^. Movement may be driven by several factors including: limited natural resources, conflicts, recurrent severe droughts, and extreme weather ^21,22^. Resource depletion in recent years has led to increased movement at longer distances in search of grazing areas and water to sustain livestock ^23,24^. *An. stephensi* has exhibited both zoophagic (common livestock species) and anthropophagic behaviors in populations in South Asia ^25^. Data on *An. stephensi* feeding preferences in Ethiopia can provide preliminary insight to the potential for extensive livestock movement as a vehicle for *An. stephensi* spread. Here we performed phylogeographic and population genetic analyses of *An. stephensi* at ten sites and used sequence barcoding approaches to evaluate hosts on which of *An. stephensi* feeds to better understand vector movement.

## METHODS

### Sample collection and site description

Sample collection was conducted in ten sites from September to November of 2018 in east and northeast Ethiopia including Semera, Bati, Awash Sebat Kilo, Gewane, Dire Dawa, Erer Gota, Degehabur, Jigjiga, Kebridehar, and Godey as previously detailed ^13^. These sites were selected based on the high prevalence of malaria infections, variation in landscape (altitude and topography), and proximity to major roads. Mosquitoes were collected using CDC light traps and pyrethrum spray collections (PSC) in houses, and larvae and pupae were sampled using the WHO dipping approach. CDC light-traps were set up from 18:00 to 6:00 local time. Pyrethrum spray collections were performed between 6:00 and 8:00 hours. Larvae and pupae were collected from suspected larval habitats including human made containers using a standard dipper (350 ml capacity) and pipettes. Adults were reared from collected larvae and morphologically-identified to species using palps, wings, abdomen, and legs based on standard identification keys ^26–28^ and confirmed molecularly through *COI* and *ITS2* sequencing. Overall, a total of 187 mosquitoes were analyzed in this study, 118 wild-caught reared from immatures and 69 wild-caught adults (Table 1).

**Table 1:**
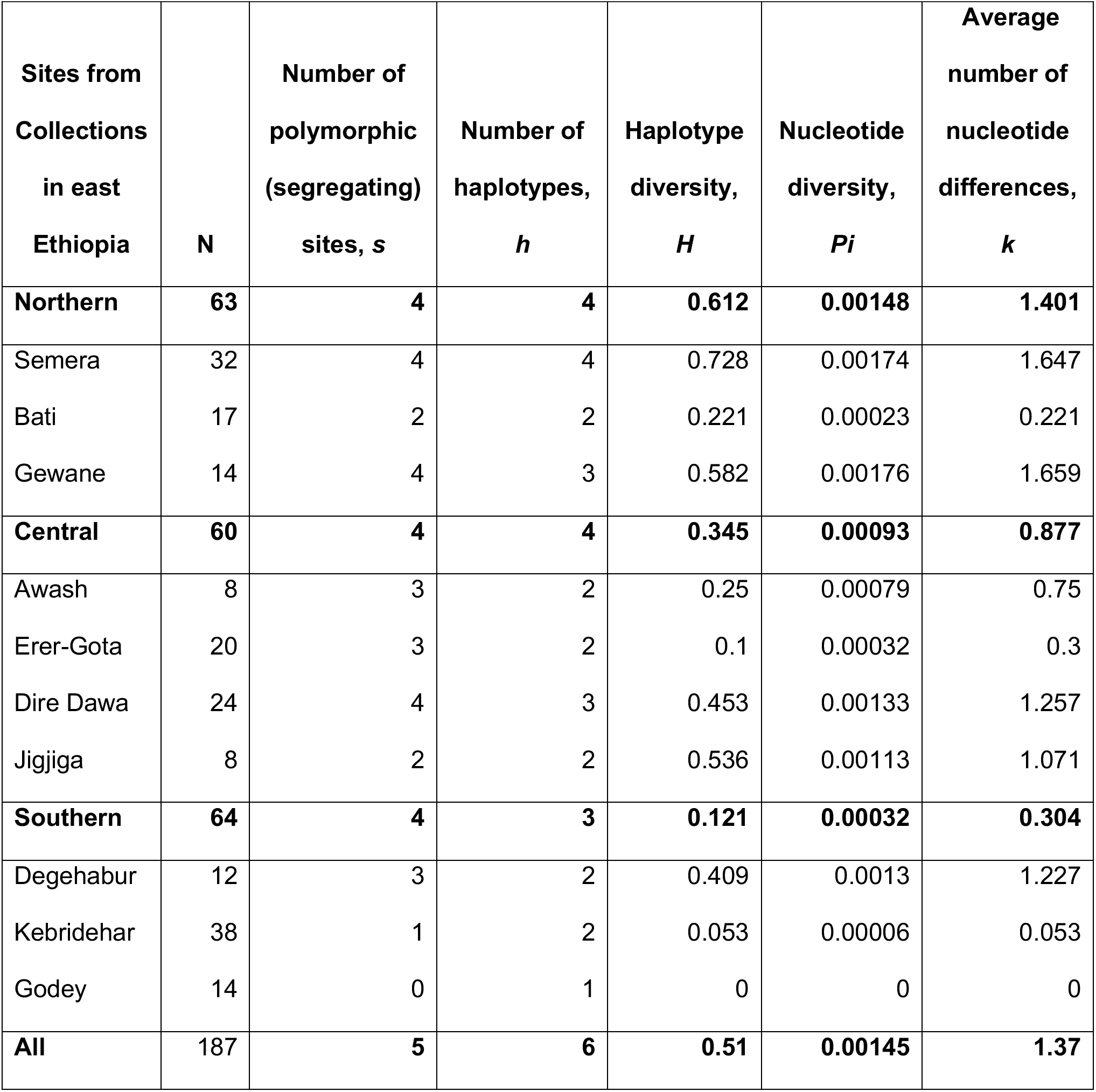
Population genetic statistics from collection sites in Ethiopia based *COI* and *CYTB* sequences

### Analysis of *An. stephensi* DNA

Two *An. stephensi* genes were selected for genetic diversity analysis. *COI* was chosen based on the availability of sequences from other countries for global phylogenetic analysis. *CytB* was chosen for its successful application in characterizing other Culicidae. While ITS2 was helpful for species identification^29^, no genetic variation was observed at this locus in these Ethiopian samples ^3^ and thus ITS2 phylogenetic analysis was not undertaken. *COI* sequences were generated as previously described ^2^. For *CytB*, two primers were designed with Primer3 software^30,31^ : CytbF 5’AGGATCTTCTACAGGACGAG3’ and CytbR 5’CATGTAGGACGAGGAGTCTA3’ and used for PCR amplification. Final reagent concentrations and components were 0.4 μM for each primer, 1X Promega GoTAQ HotStart master mix (Promega, Madison, Wisconsin), and water for a total reaction volume of 25μl. The temperature protocol was performed as follows: 94° C for 5 min, 35 cycles of 94 °C for 40 sec, 56 °C for 1 min, 72 °C for 3 min, and final extension of 72 °C for 10 min. Amplification was confirmed using visualization of a 750 bp band with gel electrophoresis. PCR products were sequenced using Sanger technology with ABI BigDyeTM terminator v3.1 chemistry (Thermofisher, Santa Clara, Ca) according to the manufacturer’s recommendations and run on a 3130 Genetic Analyzer (Thermofisher, Santa Clara, CA). *CytB* sequences were trimmed and analyzed using CodonCode (CodonCode Corporation, Centerville, MA). To confirm amplification of the correct locus, sequences were submitted as queries to the National Center for Biotechnology Information’s (NCBI) Basic Local Alignment Search Tool (BLAST) ^32^ against the nucleotide collection in NCBI’s Genbank under default parameters [max High-scoring Segment Pairs (HSP) 250, expect threshold 10, word size 28, optimized for highly similar hits, not specific to any organism].

### Population genetic analysis

#### Nucleotide Diversity Statistics

To estimate the level of diversity of the overall *An. stephensi* collection and within each site, we calculated the number of polymorphic (segregating) sites (*s*), number of haplotypes (*h),* haplotype diversity (*H)*, nucleotide diversity (*Pi*), and average number of nucleotide differences (*k*) using the program DNAsp v5 ^33^. Genetic diversity statistics were also generated for each collection site subregion designated “northern”, “central”, “southern” as listed in Table 1.

### Phylogenetic Analysis

In order to determine the evolutionary relationships between the *An. stephensi* found in Ethiopia and further elucidate the spread of the vector, we also performed phylogenetic analysis of the *COI* and concatenated *COI*+*CTYB An. stephensi* sequences generated in this and previous studies.

Alignments were created with MAFFT version 7 ^34^ and ragged ends were trimmed using Mesquite 3.51 ^35^. Phylogenetic relationships were inferred using RAxML ^36^ which is based on a maximum likelihood (ML) approach. The GTRGAMMA option that uses GTR model of nucleotide substitution with gamma model of rate of heterogeneity was applied. One thousand replicates were performed with the strategy searching for the heuristically-best-scoring tree and bootstrap analysis in one run. Best scoring trees under ML with bootstrap values from RAxML were viewed in FigTree v1.4.4 ^37^ and labeled based on geographic location.

To provide some preliminary insight into the relationship between the haplotypes and other global haplotypes, we also performed phylogenetic analysis using the *COI* sequence data available in Genbank (Supp File 1). Analyses were performed as above except that we only incorporated unique haplotypes into the final RAxML analysis. Trees were visualized in Figtree and symbols added to represent the countries.

### Minimum Spanning networks

To evaluate the frequency and relationship between Ethiopian *An. stephensi* haplotypes observed, we generated minimum spanning networks ^38^ based on the concatenated *COI*-*CytB* sequences using PopArt ^39^. Similarly, we generated a network of global populations using *COI* sequences from population data sets available in GenBank. Population data sets were only available for Sri Lanka, Pakistan, and Saudi Arabia; analysis was limited to these sets.

### Molecular blood meal analysis

DNA was extracted from abdomens of 59 wild caught adult blood fed *An. stephensi* using Qiagen DNeasy kits and used for molecular identification of bloodmeal sources. We used a sequence-based approach based on the *CytB B* gene and a universal vertebrate specific primer set to first: 1) confirm whether there was detectable vertebrate DNA present in the sample and 2) identify the vertebrate host based on querying NCBI nucleotide sequence database. This primer set can be used to distinguish a broad range of vertebrate species including human, cow, and goat^40^ (Kent and Norris 2005). The primers used were UNFOR403 5’TGAGGACAAATATCATTCTGAGG3’ and UNREV1025 5’GGTTGTCCTCCAATTCATGTTA3’.

PCR was performed using the following final reagent concentrations and components: 0.4 μM for each primer, 1X Promega GoTAQ HotStart master mix (Promega, Madison, Wisconsin), and water for a total reaction volume of 25μl. Amplification conditions were as follows: 95° C for 5 min, 35 cycles of 95 °C for 1 min, 58 °C for 1 min, 72 °C for 1 min, and final extension of 72 °C for 10 min. PCR products were confirmed using gel electrophoresis as detailed above, sequenced and compared with database sequences using BLAST for species identification.

## RESULTS

### Population genetics analysis

We were interested in the number of haplotypes that would be detected in *An. stephensi* in our study. Analysis of *COI* sequences from 187 *An. stephensi* revealed four segregating sites (polymorphisms) across the sequence that contributed to six distinct *COI*-haplotypes. All polymorphisms were synonymous mutations. Comparison with *CytB* had a single segregating site resulting in two haplotypes. Genetic diversity statistics were generated for each site including number of polymorphic (segregating) sites (*s*), number of haplotypes (*h),* haplotype diversity (*H)*, nucleotide diversity (*Pi*), and average number of nucleotide differences (*k*) Genetic diversity statistics were also generated for each collection site subregion designated “northern”, “central”, “southern” as listed in Table 1. Semera, located in the northern portion of east Ethiopia, was the most diverse (h= 4, H = 0.728) and Godey in the south was the least diverse (1 haplotype, H=0). We observed a trend of declining diversity from north to south (Figure.1), with northern sites having the greatest level of diversity (h=5, H=0.63) and the southern sites with the least (h=3, H= 0.121).

**Figure 1.**
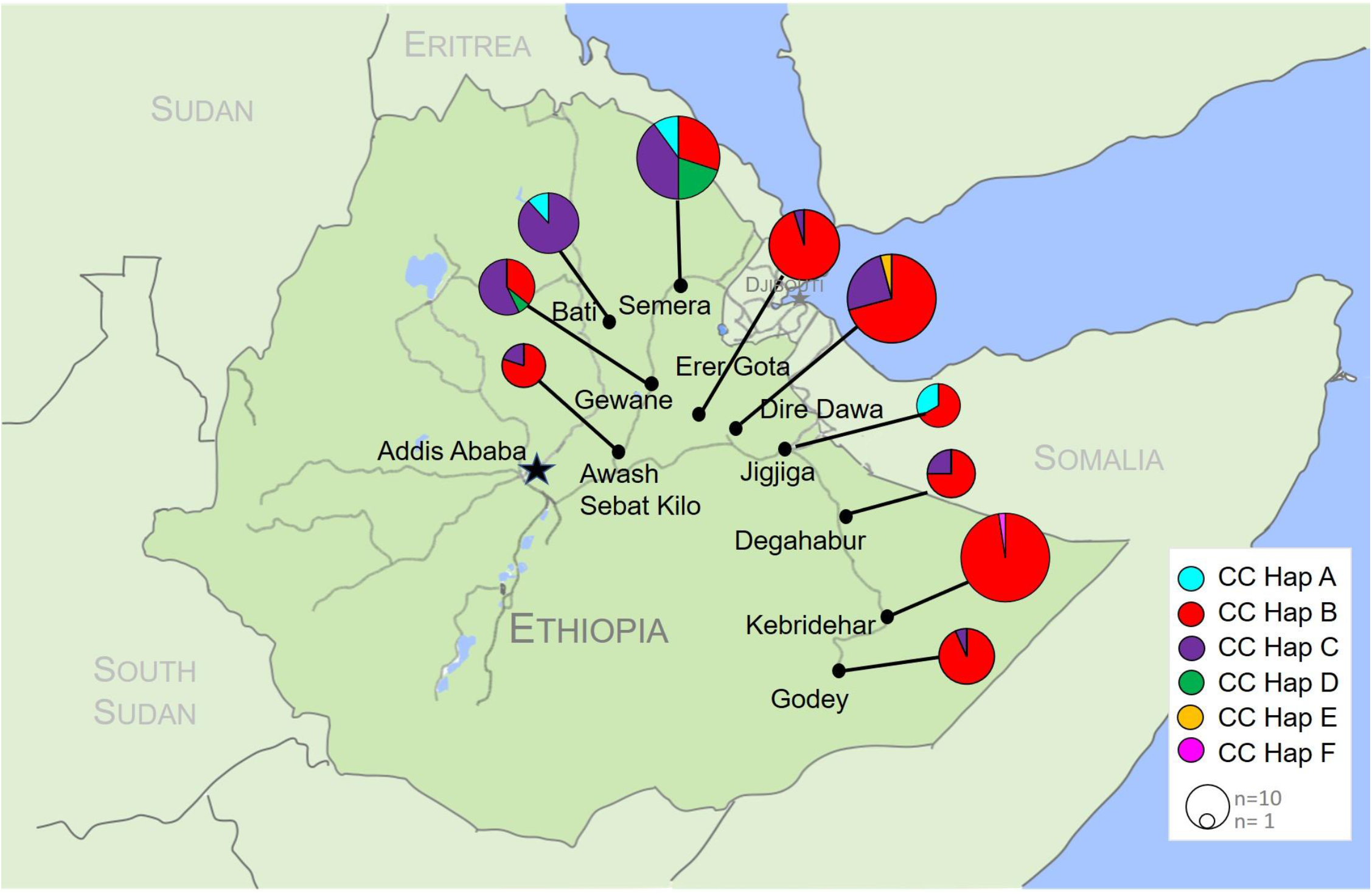
Distribution of *An. stephensi COI-CYTB* (CC) haplotypes (Hap) per site in east Ethiopia, Nov – December 2018. Pie charts are proportional to the sample size. Each color represents a different haplotype.

### *An. stephensi* appear geographically differentiated

In addition to different levels of variation in the northern compared with the southern sites, haplotypes were distributed differently across the three designated subregions. The most prevalent haplotype in the collection was CC-Hap B and it was observed in all three subregions. CC-Hap B was also the predominant haplotype in the southern and central subregions (93.8% and 82.9% respectively). However, in the northern subregion, where the highest number of distinct haplotypes was observed, the predominant haplotype was CC-Hap C (55.6%). Phylogenetic analysis supports population differentiation between the northern and southern sites with bootstraps >90 for three major clades that carry haplotypes that vary in frequency across these subregions (Fig. 2a). We also constructed a minimum spanning networking based on the *COI-CytB* haplotypes to further evaluate the relationship between the haplotypes (Fig. 2b). The most central nodes (representing specific haplotypes) were CC-Hap A and CC-Hap B. Each haplotype in the network differed by 1-2 nucleotides.

**Figure 2.**
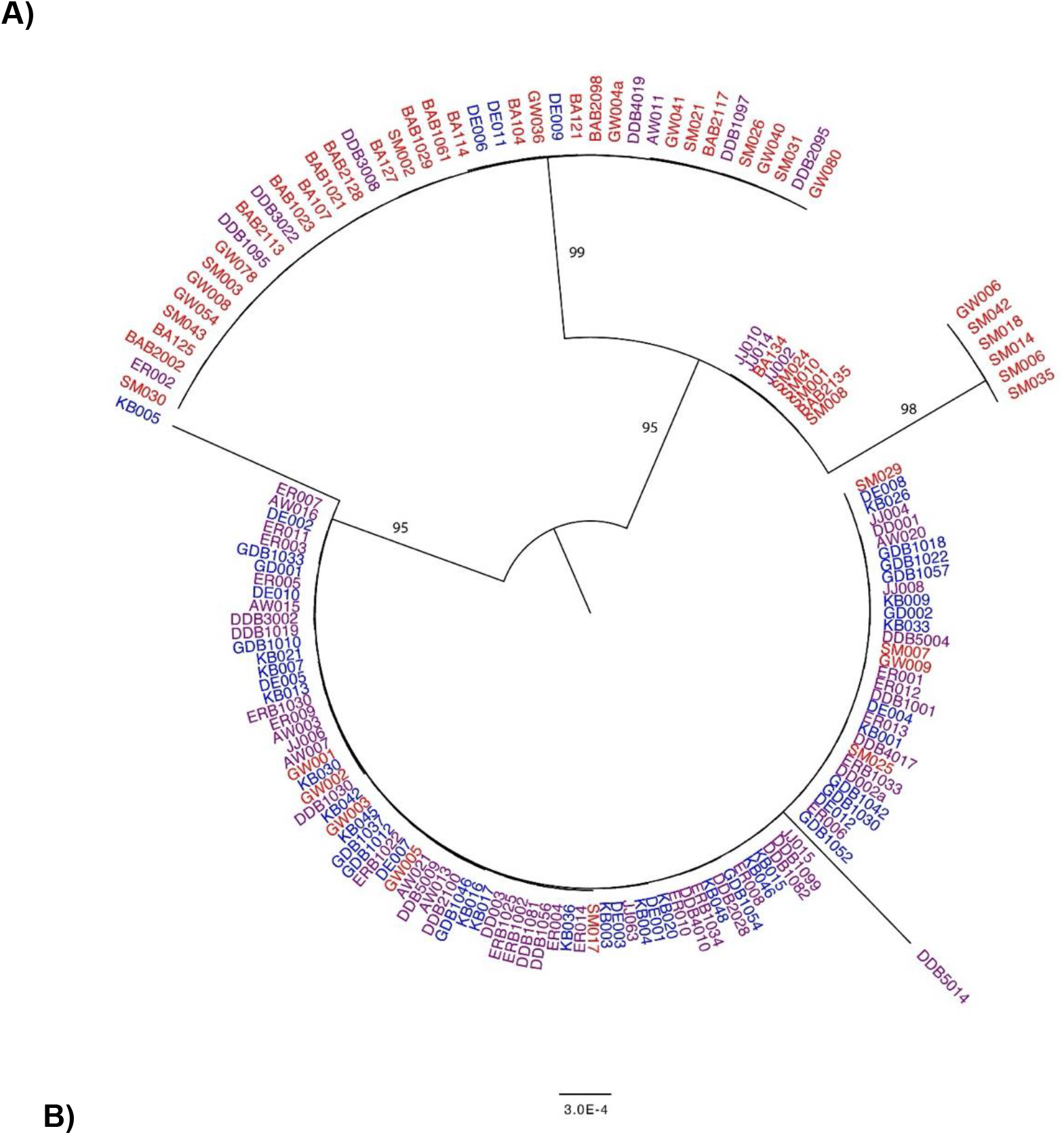
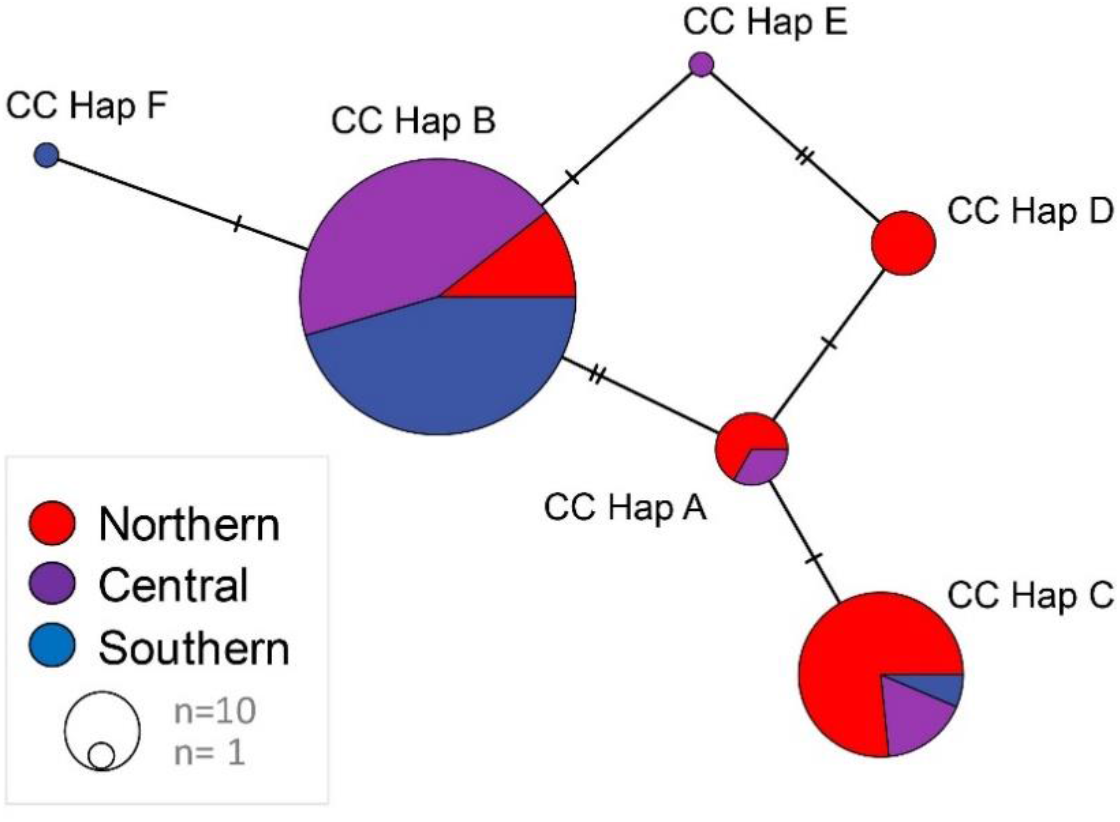
Relationship between *An. stephensi COI-CYTB* sequences in Ethiopia. Colors represent sub-regions with east Ethiopia collections. **A)** Phylogenetic tree of *An. stephensi COI-CTYB* sequences. Only bootstrap values > 70 are shown. **B)** Minimum spanning networking of Ethiopian *An. stephensi COI-CYTB* (CC) haplotypes. Each node represents a haplotype and the proportion of that haplotype contributed by each Ethiopian region. The size of the nodes is proportional to the sample size. The ticks between nodes represent the number of nucleotide differences.

### Ethiopian *An. stephensi* share haplotypes with several countries

*CytB* sequences were only available for three countries so only *COI* sequences were used for global diversity comparisons. We compared the *COI* sequences from Ethiopian *An. stephensi* sequences to the sequences available in the NCBI’s Genbank representing countries in South Asia (India, Pakistan, Sri Lanka), the Middle East including Iran and the Arabian Peninsula (Saudi Arabia, United Arabic Emirates), and the Horn of Africa (Djibouti, Ethiopia). Of the six Ethiopian *COI* haplotypes detected, two were observed in other countries, designated *COI*-Hap1 (all CC-Hap A) and *COI*-Hap2 (all CC-Hap B). The minimum spanning network of the available population *COI* sequence datasets revealed *COI*-Hap1 was the most central node, with the most proximal nodes representing country-specific haplotypes for Ethiopia, Sri Lanka, and Pakistan that differ from the central node by a single nucleotide (Fig 3). The haplotype represented by the central node was also the predominant haplotype in Pakistan (26/28). Phylogenetic analysis of the global *COI* haplotypes reveals that *COI*-Hap1 is the most widely dispersed haplotype, observed in most sites including Djibouti and other locations in the Arabian Peninsula, South Asia, and the Middle East (Fig 4). Saudi Arabia, with the most basal sequences and strong differentiation from the other *An. stephensi* sequences (bootstrap = 90), lacked the *COI*-Hap3 haplotype and did not share any haplotypes with Ethiopia or any of the other countries (Fig 3, Fig 4).

**Figure 3.**
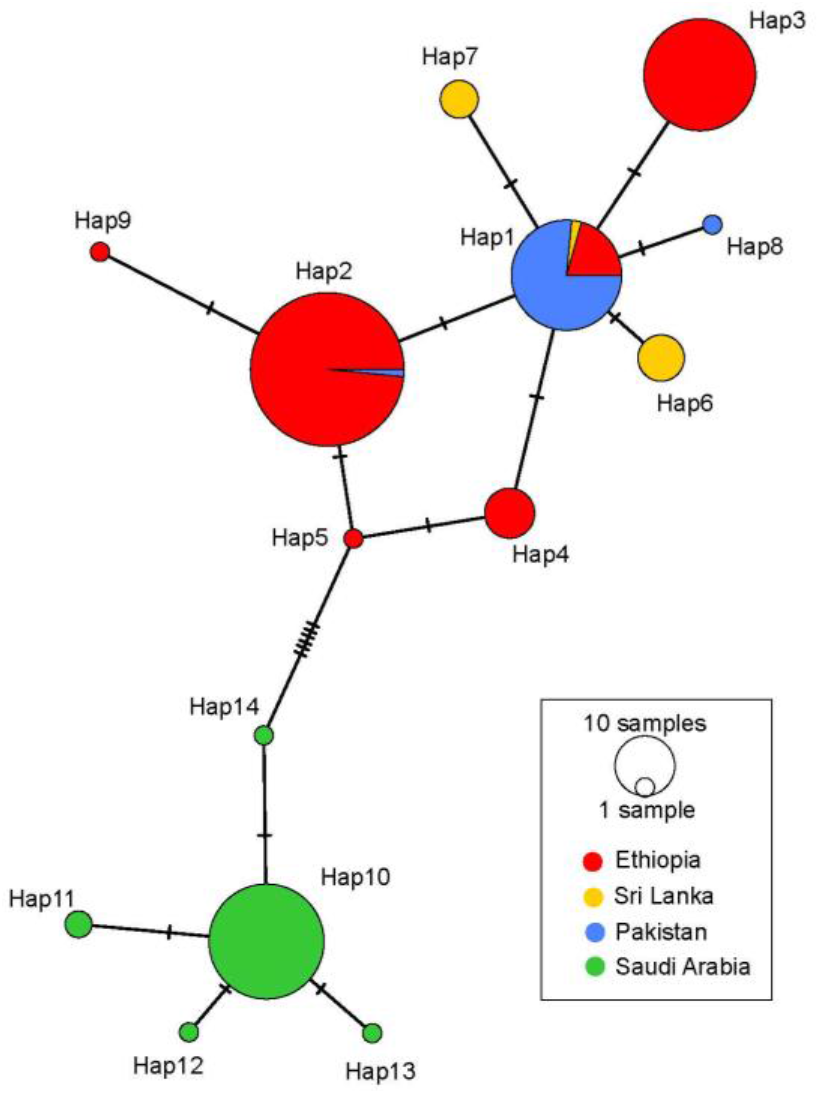
Minimum spanning network of *An. stephensi COI* haplotypes (H) from Ethiopia, Sri Lanka, Pakistan and Saudi Arabia. Each node represents a haplotype and the proportion of that haplotype contributed by each country. The size of the nodes is proportional to the sample size. The ticks between nodes represent the number of nucleotide differences.

**Figure 4.**
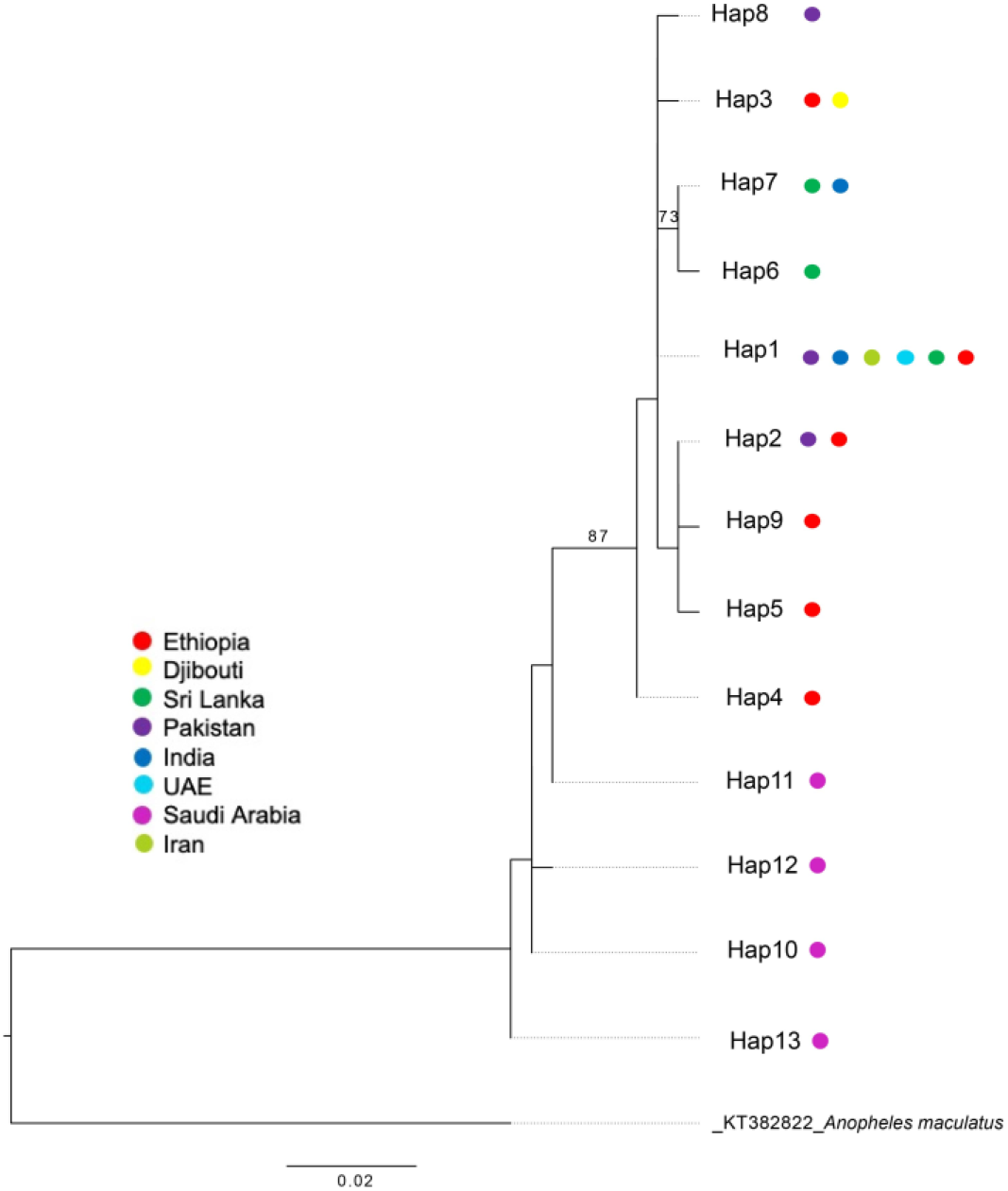
Global *Anopheles stephensi* COI haplotype tree. Each dot represents the country where the COI haplotype had been observed. Only bootstrap values > 70 are shown.

### *An. stephensi* in East Ethiopia exhibit zoophilic feeding behaviors

Given the evidence of a recent spread of *An. stephensi* in East Ethiopia, we were very interested in whether the *An. stephensi* exhibited zoophilic feeding behaviors that may encourage its range expansion through livestock movement. Of the 59 wild-caught adult *Anopheles stephensi* mosquitoes with available DNA from abdomens, 36 (61.0%) were confirmed to be blood-fed based on the presence/absence PCR assay for vertebrate DNA. These originated in Semera (n=27), Erer Gota (n=4), Gewane (n=2), Kebridehar (n=2) and Godey (1). Of the 36 blood-fed *An. stephensi*, 75.0 % (27/36) had vertebrate DNA identifiable with sequencing. Of the 27 bloodmeals, 22 were from goat, three from cow, one from human, and one from dog. Of the 22 samples confirmed to have fed on goat, 68.2% (15/22) showed signs of intraspecies mixed feeding based on the presence of double peaks in sequencing data. Of the 9 samples considered blood-fed but unidentifiable, four samples showed evidence of cross species feedings that could not be resolved by sequence analysis and five samples presented positive presence/absence PCR results for blood feeding but produced unsuccessful sequencing results. Overall, we found evidence of both livestock and human feeding.

## DISCUSSION

### Population genetic analysis reveals signatures of a range expansion

Our analyses revealed multiple mitochondrial gene haplotypes in *An. stephensi* populations in Ethiopia and some geographic differentiation. In addition, a general decline in genetic diversity from the most northern site, Semera, to the most southern, Godey, was observed. The presence of geographic differentiation supports *An. stephensi* populations in Ethiopia being relatively new as extensive migration would have been expected to lead to more similar haplotype frequencies across the regions. A cline in genetic diversity supports expectations for recent range expansion, where the most recent population would have the lowest level of diversity ^41–43^. In this study, the observation of higher diversity in the northern sites and lower diversity in the southern sites suggests that *An. stephensi* in the south represent a more recent introduction relative to the population in the north.

### Global comparison reveals continental origin of *An. stephensi* in Ethiopia

Phylogenetic and network analysis (Fig 3, 4) revealed that both the *An. stephensi* in Ethiopia and Djibouti are more closely related to *An. stephensi* from South Asia than those from the Arabian Peninsula (primarily Saudi Arabia). These results support South Asia as the place of origin for *An. stephensi* in the Horn of Africa. Of the countries included in this study, Pakistan is the strongest candidate for country of origin of the *An. stephensi* populations in the HOA, based on the centrality of the predominant *COI*-Hap1 haplotype observed in Pakistan relative to the haplotypes found in recent populations from Ethiopia and Sri Lanka. However, conclusions about the specific origin (i.e. country) require additional sampling and sequencing from long-standing *An. stephensi* populations from countries not well represented in the available database including Sudan, Afghanistan, Yemen, Oman, and China.

### Haplotype geographic distributions support multiple introductions into Ethiopia

It is also possible that there were multiple introductions of *An. stephensi* into Ethiopia given that there were multiple haplotypes that emerged from the central *COI*-Hap 1 and that secondary haplotypes appear geographically structured. If the *COI*-Hap1 represents the haplotype of origin, then the candidate sites for introduction would be those where this haplotype is observed. In our analysis, we see the central *COI*-Hap1 haplotype in Semera and Bati in the northern subregion and Jigjiga in the central subregion (Fig. 1). Based on these data, it is likely *An. stephensi* was introduced into the north and/ or central subregions of eastern Ethiopia and subsequently spread into the south. The proximity of these sites to Djibouti highlights the importance of additional molecular surveillance in that country to understand the impact of movement between Djibouti and Ethiopia on the introduction of *An. stephensi* in Ethiopia.

### Potential factors for spreading into the Horn of Africa and throughout Ethiopia

The geographic distribution of *An. stephensi* genetic diversity provides insight into potential modes of introduction and spread in the Horn of Africa. The southern populations are likely the most recent populations and the central “origin” *COI*-Hap 1 haplotype is found in sites most proximal to Djibouti. These three sites (Semera, Bati, and Jigjiga) are located on major roads that lead into and out of Djibouti. These roads may be central to the movement of *An. stephensi* from Djibouti to Ethiopia. Particularly, travel by vehicle compartments related to the transport of goods may have facilitated the introduction. Most of our sample sites are proximal to major roads, therefore, additional sampling of more remote regions off the major roads for comparison can further elucidate the role of the roads in the movement and transport of *An. stephensi*. Several of these sites have an international airport and introduction via aircraft is also possible.

Additional factors may be contributing to the ongoing spread of *An. stephensi* into other parts of the country. Pastoralist activity has been proposed as a potential factor particularly in the south east portion of Ethiopia. This movement, often in search of rangeland includes the potential movement of water containers which may hold *An. stephensi* eggs. Whether the long-distance transport of livestock could also lead to the transportation of the vector is not yet clear. Given the evidence for both zoophilic and anthropophilic feeding, *An. stephensi* could be attracted to these mobile herds and thus might be transported long distances in this way. Previous reports show that *An. stephensi* may switch between anthropophagy and zoophagy depending on the presence of livestock in south Asia 44.

Climate variables may also play a role in the movement of *An. stephensi* and ultimately the pattern of population structure. Factors such as average rain fall, temperature, predominant winds and wind speed may influence the movement of mosquitoes or induce local adaption for rapid expansion. While Jigjiga and Semera are predicted suitable habitats for *An. stephensi*, many of the other sites are not, including Degehabur, Kebridehar, and Godey ^45^. *An. stephensi* has been detected as far south as Godey (1200 km from Semera and 400km from Jigjiga), which has very low levels of rain fall relative to some of the northern sites. The fact that *An. stephensi* breeds in man-made containers suggests that the movement of human populations may be a more relevant factor for the distribution and population structure of *An. stephensi* relative to other *Anopheles* spp. Also, questions remain about how these *An. stephensi* populations may have adapted to thrive in this environment.

Additional multilocus or genomic analyses from additional time points could provide important information on the relative geneflow across East Ethiopia to compare with the factors listed above. Because we see some predominantly northern haplotypes in the south and some predominantly southern haplotypes in the north, there may have been some *An. stephensi* migration between the two subregions. Analysis is ongoing to quantify the degree of movement between the sites studied here.

## CONCLUSION

Our data on Ethiopian *An. stephensi* genetic diversity suggest a South Asia origin of the Horn of Africa *An. stephensi* and a variable timeline of introduction between north/central vs. south east Ethiopia. Future studies with expanded genomic analysis will further inform our understanding of migration rates, the role local adaption on the spread of *An. stephensi* into and throughout the Horn of Africa. Collaboration across countries where this mosquito is well established and countries where it is emerging can facilitate a complete understanding of the spread of *An. stephensi* into Africa.

## Acknowledgements

This research was funded by a NIH Research Enhancement Award (1R15AI151766) and Baylor University. In addition, the US President’s Malaria Initiative provided funding to PMI VectorLink (Abt Associates) for the training, collection, and identification of mosquitoes. SI was funded by PMI. The findings and conclusions in this report are those of the author(s) and do not necessarily represent the official position of the Centers for Disease Control and Prevention”

## Contributions

TEC, SRI, SZ, DJ, and MB contributed to the conception and design of the project. SI and MB organized/lead the collection of specimens. TEC, JS, SHC, JNS, KL generated the data. TEC, JS, and SHC analyzed the data. TEC, SY, DG, DJ, SRI, SZ, MB contributed to the writing of the paper. All authors read and approved the final manuscript.

## Data availability

Data used in this study are included as supplemental files and/or will be available in National Center for Biotechnology Information Nucleotide Database.

## Ethics declarations

Wild mosquitoes used for this study were collected from dwellings and animal houses, following homeowners’ verbal consent. The study protocol for the collection was reviewed by the Centers for Disease Control and Prevention, USA, and determined to be research, not involving human subjects (2017-227).

